# Selection on a pleiotropic color gene block underpins early differentiation between two warbler species

**DOI:** 10.1101/853390

**Authors:** Silu Wang, Sievert Rohwer, Devin R. de Zwaan, David P. L Toews, Irby J. Lovette, Jacqueline Mackenzie, Darren E. Irwin

**Affiliations:** Department of Zoology, 6270 University Blvd, University of British Columbia, BC, Vancouver, BC, V6T1Z4, Canada; Department of Biology and Burke Museum, Box 353010, University of Washington, Seattle, WA, 98195, USA; Department of Forest and Conservation Sciences, 2424 Main Mall, University of British Columbia, BC, V6T1Z4, Canada; Department of Biology, 619 Mueller Laboratory, Pennsylvania State University, University Park, PA, 16802, USA; Fuller Evolutionary Biology Program, Cornell Lab of Ornithology, 159 Sapsucker Woods Road, Ithaca, NY, 14850, USA

**Keywords:** speciation, admixture mapping, cline, plumage, RALY, ASIP, hybrid zone, Setophaga

## Abstract

When one species gradually splits into two, divergent selection on specific traits can cause peaks of differentiation in the genomic regions encoding those traits. Whether speciation is initiated by strong selection on a few genomic regions with large effects or by more diffused selection on many regions with small effects remains controversial. Differentiated phenotypes between differentiating lineages are commonly involved in reproductive isolation, thus their genetic underpinnings are key to the genomics architecture of speciation. When two species hybridize, recombination over multiple generations can help reveal the genetic regions responsible for the differentiated phenotypes against a genomic background that has been homogenized via backcrossing and introgression. We used admixture mapping to investigate genomic differentiation and the genetic basis of differentiated plumage features (relative melanin and carotenoid pigment) between hybridizing sister species in the early stage of speciation: Townsend’s (*Setophaga townsendi*) and Hermit warblers (*S. occidentalis*). We found a few narrow and dispersed divergent regions between allopatric parental populations, consistent with the ‘divergence with gene flow’ model of speciation. One of the divergent peaks involves three genes known to affect pigmentation: ASIP, EIF2S2, and RALY (the ASIP-RALY gene block). After controlling for population substructure, we found that a single nucleotide polymorphism (SNP) inside the intron of RALY displays a strong pleiotropic association with cheek, crown, and breast coloration. In addition, we detect selection on the ASIP-RALY gene block, as the geographic cline of the RALY marker of this gene block has remained narrower than the plumage cline, which remained narrower than expected under neutral diffusion over two decades. Despite extensive gene flow between these species across much of the genome, the selection on ASIP-RALY gene block maintains stable genotypic and plumage difference between species allowing further differentiation to accumulate via linkage to its flanking genetic region or linkage-disequilibrium genome-wide.

## Introduction

Examining the genomic distribution of differentiation between two populations can reveal targets of divergent selection, advancing our understanding of the speciation process [1–6]. At the onset of divergence between sister taxa, differentiation at narrow regions of chromosomes, or “islands of differentiation”, is expected [7–12]. Strong selection on a few genes of large phenotypic effects can result in a small number of highly differentiated regions. Physical linkage results in elevated divergence of the nearby neutral regions, a phenomenon known as divergence hitchhiking [2,7,11]. More differentiation can accumulate as speciation progresses, thus extending regions of high differentiation and forming “continents of divergence” [4]. Alternatively, speciation can be initiated via many regions of small effects on traits under multifarious selection (one selection targeting multiple traits) and gradually accumulate genome-wide differentiation via correlated differentiation across the genome [13–18]. A central debate in speciation research has therefore been whether speciation occurs through gradual differentiation throughout the genome or more rapidly at a few key regions. Genomic analyses of sister species in the early stages of speciation uncovers the evolutionary process of speciation, thus inform this debate.

Islands of divergence may be associated with divergent (and often diagnostic) traits that are involved in reproductive isolation [19–21]. Divergent phenotypic traits frequently play important roles in social interactions at geographic boundaries between closely-related species [22, 23]. These features are commonly involved in mate-choice [19, 20] or aggressive signaling [21, 24] that are divergently selected [25–30] and facilitate species recognition and reproductive isolation [31, 32]. We are just beginning to understand the genetic underpinnings of traits related to species divergence [26,27,29,33,34]. Integrating the genetic basis underlying pigmentation with corresponding evolutionary forces will deepen our understanding of the formation and maintenance of species barriers.

Carotenoid and melanin pigmentation are commonly involved in species-diagnostic traits in avian signals [19,25–28,32,35]. At the boundaries between closely-related species, the intensity of coloration might not differ, but rather the differential patterning of the colors is involved in mate choice favoring conspecifics [32,35,36]. In such cases, the regulatory genes for melanin and carotenoid pathways are strong candidates for differentiation between species as these loci are likely to cause differences in plumage pattern.

Hybrid zones, locations where divergent lineages interbreed, provide opportunities to understand the relationship between species-diagnostic features and genomic differentiation. Social signals can be crucial for conspecific mate recognition and intrasexual competition for mating opportunities [19,20,37–39]. When the plumage signals established in isolated populations come into secondary contact, hybrids can be selected against due to their intermediate or mismatched parental signals interfering with mate recognition and competition [37, 38].

The hybrid zone between *Setophaga townsendi* (abbreviated as *townsendi*) and *S. occidentalis* (abbreviated as *occidentalis*) along the Cascade mountains harbors extensive gene flow between these closely related species that started diverging an estimated 400,000 years ago [40–44]. Males of this species pair differ in several plumage features related to both carotenoid and melanin patterning on the crown, cheek, breast, and back. For example, *townsendi* has a melanin-based cheek patch (Figure 1A right), whereas *occidentalis* displays a completely carotenoid-based cheek (Figure 1A, left). Because apparent hybrids predominantly resemble *occidentalis* in crown and cheek coloration (Figure 1A center), with intermediate breast (the extent of yellow plumage) and back colors (green plumage in the mantle), Rohwer & Wood [43] predicted that face coloration of Hermit Warblers and hybrids would be controlled by a single-locus dominant allele. In addition, whether the other carotenoid and melanin patterning differences between species are underpinned by the same genetic mechanism is an open question. These signals can be important in male-male competition for territories, as individuals in the hybrid zone demonstrate an aggression bias towards different plumage types [42, 45].

**Figure 1.**
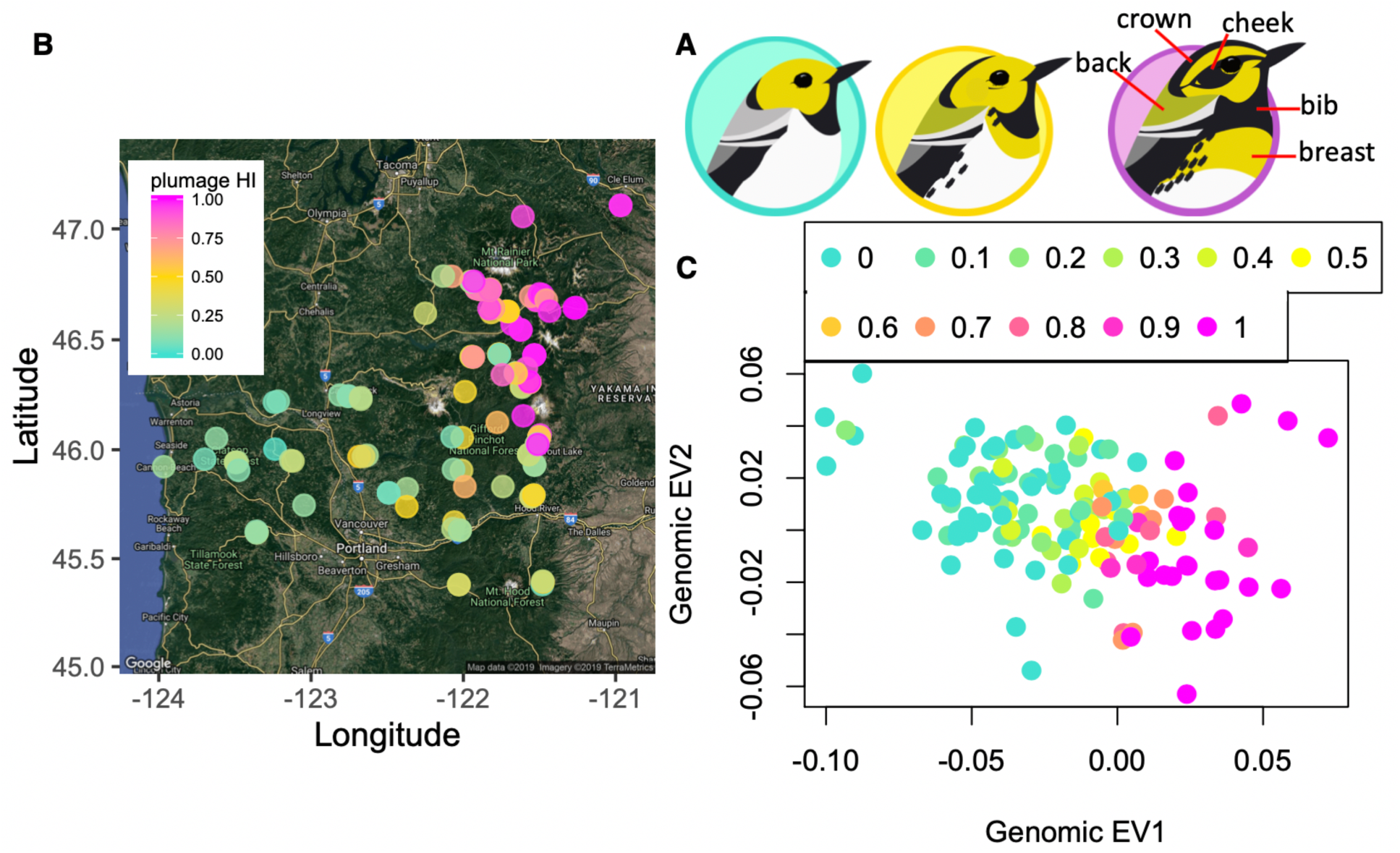
**A**, illustration highlighting the key plumage difference between *occidentalis* (left, turquoise), hybrids (center, yellow), and *townsendi* (right, magenta) (illustration by Gil Jorge Barros Henriques). **B**, a map showing site mean plumage hybrid index based on 8 plumage traits (0 for pure *occidentalis*, in turquoise; 1 for pure *townsendi*, in magenta). **C**, Genomic eigenvector 1 (EV1) and eigenvector 2 (EV2) with individual datapoints colored by plumage hybrid index. The genomic EV1 reflects the variation among individuals that are *occidentalis*-like (low genomic eigenvector EV1) versus *townsendi*-like (high genomic EV1).

We investigated whether speciation in this young species pair occurred through gradual widespread differentiation across the genome or via a small number of restricted regions. If selection for coloration phenotypes has targeted a small number of specific genomic regions, genomic islands of differentiation should be narrow, dispersed, and associated with loci involved in pigmentation pathways [3,7,8]. In addition, we asked the following questions regarding the genetic basis of species-specific traits: (1) is there an island of divergence pleiotropically underpins species-specific features (crown and cheek darkening, breast yellow, bib size, greenish back); (2) Is cheek darkening influenced largely by an allele of dominant effect, consistent with the prediction made two decades ago by Rohwer & Wood (1998); and (3) Is there stable selection maintaining differentiation in the gene region underlying key species diagnostic features?

## Materials and Methods

### Sampling

Whole specimens of *Setophaga* warblers were collected over two historical sampling sessions in the Washington Cascade hybrid zone [41, 43]. The first sampling (*N* = 314 individuals; 35 sites) was carried out in 1987-1994 [43], while the second (*N* = 127; 11 sites) was in 2005-2008 [41] and covered a subset of the sites from the original sampling. We accessed these specimens at the Burke Museum of Natural History and Culture (University of Washington, Seattle, Washington).

We carried out a third round of sampling using a catch-and-release approach during the breeding season (early May to mid-July) in 2015-16 (Figure 1B). Upon locating a territorial male by song, a mist net with a playback (of a locally recorded song) at the bottom was set up nearby. After capturing an adult, photographs and a blood sample were taken for further analysis. We re-sampled the sites that were sampled in 1986-94 [43]. For details, see previous work in this system [46].

### Plumage measurements

Melanin- and carotenoid-based plumage traits allow identification of the two species (Figure 1B), but there is also variation within each species [43, 47]. To quantify plumage variation within and between populations, we focused on five distinct plumage traits in males: the relative amount of black (melanin) and yellow (carotenoid) in the 1) cheek, 2) crown, and 3) breast and the 4) throat bib, as well as, 5) the intensity of green chroma on the back (Figure 1B).

For each of the 242 warblers captured in the field, we took three pictures from different angles: 1) frontal with head tilted up (for bib and breast measurements), 2) profile (cheek), and 3) from above (crown) (Figure S1). Plumage color metrics were measured using Adobe Photoshop CC in CIE (*Comission Internationale de l’Eclairage*) LAB color space. LAB color space is a 3-dimensional space consisting of 3 distinct, perpendicular axes: 1) Luminosity (L) ranging from 0 (black) to 100 (white), 2) ‘a’ ranging from green (negative) to red (positive), and 3) ‘b’ ranging from blue (negative) to yellow (positive; Adobe 2017). We chose this color space because it linearizes the variables of interest along three distinct axes: black (‘L’), yellow (along the ‘b’ axis), and green (‘a’).

For the cheek and mantle, we selected a standardized area and averaged the pixels to record the ‘b’ and ‘a’ values, respectively. For the cheek, we selected and averaged the entire area from above the eye (but excluding the eye) to the throat badge, and from the base of the bill to the mantle using the profile photos. Differences in ambient light conditions at the time a picture is taken can confound comparison of color metrics among individuals. To address this, we used the white-balance feature in Photoshop, using the white plumage of each individual’s belly as a standard, to correct for differences in ambient light among photos and standardize the color metrics. We acknowledge that without spectral analysis, we do not incorporate UV reflectance which is a ubiquitous aspect of signaling in avian systems [48]. However, our methods allow us to estimate the relative intensity of melanin- and carotenoid-based plumage traits during the breeding season.

To measure the size of the black bib, we used the program Analyzing Digital Images (ADI;[49]). A scale was included in all photos to standardize size measurements among individuals. We measured bib size (Figure 1A) by creating a polygon around the bib and calculating the area (± 0.1 mm^2^).

### GBS pipeline

The GBS pipeline details have been described in [46]. Following [50], we prepared genotyping-by-sequencing (GBS) [51] libraries from individual DNA samples. In brief, genomes were digested with restriction enzyme and ligated with barcode and adaptors, amplified with PCR, and size selected (fragment length of 300 - 400 bp) for sequencing. Libraries were sequenced at Genome Quebec with paired-end sequencing (read length = 125 bp) on an Illumina HiSeq 2500 automated sequencer. The resulting sequences were processed following a GBS pipeline [52]. All the sequence can be acquired through GenBank (accession number: PRJNA573930, ID: 573930). We demultiplexed the reads with a custom script and trimmed them using Trimmomatic 0.36 (Bolger et al 2014) [TRAILING:3 SLIDINGWINDOW:4:10 MINLEN:30], then we aligned reads to a *Taeniopygia guttata* reference version 3.2.4 [53] using bwa (Li et al 2009) (default settings). We assumed synteny of *Setophaga* and *T. guttata* genomes given the limited rearrangement in avian genomes [54, 55], and the conclusions of this study would be unlikely to be affected by a moderate number of rearrangements. We conducted SNP calling with GATK (McKenna et al. 2010), which produced 4,097,089 SNPs. The SNP filtering was done with VCFtools [56], which includes removing indels, genotype quality (GQ) > 20, minor allele frequency (MAF) ≥ 0.05, removing loci with >30% missing data, and only including biallelic SNPs, resulting in 21,852 SNPs remaining.

### Whole Genome Sequencing (WGS)

In addition, we selected 5 samples from *occidentalis* in Pinehurst, CA, U.S.A. (UWBM 66152, 66153, 66148-66150) and 5 samples from inland *townsendi* in Tok, AK, U.S.A. (UWBM 84816-84819, 84860) from the Burke museum for whole genome re-sequencing. For DNA extraction, we used 2 mm^3^ of tissue digested in Qiagen buffers—following the tissue extraction procedure—and separated DNA using UPrep spin columns (Genesee). We standardized DNA concentrations with after quantifying concentrations with a Qubit fluorometer, and then generated sequencing libraries with the llumina TruSeq Nano kit—which includes an 8-cycle PCR enrichment—selecting 350 bp insert sizes. We individually indexed each sample, and sequenced the combined libraries across a single lane of an Illumina NextSeq using the paired-end 150 bp sequencing chemistry. We combined these 10 samples with other wood warblers from other projects, but consistently included 24 individuals per sequencing lane in order to generate comparable coverage across samples.

### Genomic differentiation

To quantify the level of differentiation throughout the genome, we calculated *F*_ST_ [57] with VCFtools [56] between allopatric *townsendi* (Montana and Idaho, USA; *N* = 38) and *occidentalis* (Oregon and California, USA; *N* = 23) for each of the filtered SNPs.

The genomic architecture of differentiation from the GBS was compared to that of the WGS data. The reads from the WGS data were aligned to the same *Taeniopygia guttata* reference (version 3.2.4) [53] with bwa (Li et al 2009) (default settings). ANGSD [58] was employed to calculate genotype frequencies accounting for genotyping uncertainty before estimating *F_ST_* for each of the non-overlapping (step size = 10kb) 10kb windows.

### Admixture mapping

To identify SNPs that are associated with variation in the five plumage traits, we conducted admixture mapping with GenABEL package in R [59] for the subset (N = 242) of the GBS data produced in [46] that had pigment quantification (see above): 189 birds sampled from hybrid zone (around the center of the hybrid zone in Washington, U.S.A.), 46 from the *occidentalis* zone (Southern Oregon and California, U.S.A.), and 7 from the *townsendi* zone (Northeast Washington & Montana, U.S.A.). The GenABEL function *ibs*, which calculates identity-by-state relatedness among individuals, was used to control for population structure. Genomic control of inflation factor λ (which represents the effect of genetic structure and sample size) was conducted [59]. Briefly, λ was estimated from the genomic data assuming that randomly-selected markers from the genome are not associated with the trait (after controlling for population substructure). λ was then used to correct the test statistic χ^2^ of each association test, so that the test is relative to the null hypothesis of no phenotype-genotype association [59]. Because the cheek darkening and breast yellow intensity were left-skewed, we rank order transformed these phenotype data.

To find the number of independent hypotheses for multiple hypothesis correction, we estimated the total number of independent linkage blocks (a proxy for the number of independent hypotheses) in the dataset. Principle Component Analysis calculating genotypic covariance [60, 61] was run until 99.5% of the variance was explained with the R package SNPRelate [62], producing 210 PCs. We applied a Bonferroni correction to the independent blocks, such that significance level alpha = 0.05/210. Because we conducted admixture mapping on five plumage traits, we further applied Bonferroni correction across the 5 plumage traits, which gave rise to the final alpha value of 0.05/(210×5) ≈ 4.76×10^-5^. Candidate SNPs (i.e., those with *p*-values below this alpha value) were annotated in Ensembl Zebra Finch (taeGut3.2.4;[63]).

### Geographical cline analysis on candidate loci

As one candidate gene, RALY (see Results), stood out in the above analysis as particularly strongly associated with plumage variation, we investigated the spatial and temporal variation in this locus relative to the plumage hybrid index and rest of the genome (using samples described previously [46]). We fit the relationship between RALY allele frequency and location using an equilibrium geographic cline model [64, 65].

Geographical cline analysis followed [46]. Briefly, we collapsed the two-dimensional sampling into a one-dimensional transect by measuring the location of each site to the 0.5 isocline of the plumage hybrid index (HI) in the historical sampling [43], as follows. First, a Local Polynomial Regression Fitting (LOESS) model in R [66] was used to fit variation in HI across the hybrid zone, and this model was used to estimate the HI = 0.5 isocline in the 1987-94 sampling [43]. Then for each site, the shortest distance to the 0.5 isocline was calculated with ‘sp’ package [67]. The sites east or west of the isocline were specified as having positive versus negative distance values, respectively. We added 1200 km to the distance score of each site so that all distance values are above zero, while the relative distance of each site to the isocline is preserved. Then the data was fit to the equilibrium sigmoidal cline model [64, 65] 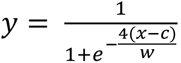, in which cline center (*c*) and width (*w*) was estimated (where *y* is HI and *x* is location with respect to the HI = 0.5 isocline). To examine whether selection (i.e., divergent selection and/or selection against hybrids) is acting on the candidate loci, we followed [46] and tested whether the increase in cline width (*w^2^_2015-16_* -*w^2^_1987-94_*) is significantly less than expected under the neutral diffusion model [68]. The change of *w^2^* (between sampling periods) as opposed to *w* allows comparison of the observed change in cline width to the neutral expectation [46]. To understand the RALY cline width change relative to the plumage and genomic cline, we compared *w^2^_2015-16_* -*w^2^_1987-94_* relative to the plumage and genomic HI cline [46], which was respectively based on the scores of the 8 plumage landmarks (with 0 representing pure *occidentalis* and 1 being pure *townsendi*), and the scaled genomic PC1 (with 0 representing pure *occidentalis* and 1 being pure *townsendi*).

## Results

Across the Cascade mountain range, the *occidentalis* plumage type was observed on the southwest and the *townsendi* plumage type was on the northeast [43, 46] (Figure 1 B). The overall pattern of genomic differentiation was consistent with the plumage divergence: *occidentalis* and *townsendi* plumage types were most different along genomic PC1. There was low genome-wide weighted average *F*_ST_ [57] between allopatric *occidentalis* and *townsendi* populations (Weir and Cockerham’s *F*_ST_ = 0.03; Figure 2A). However, four high regions of differentiation (*F*_ST_ > 0.6) were found (Figure 2A, Figure S2, gene association and functions summarized in Table S2) that map to Zebra Finch chromosome (chr) 5 (nucleotide position 25064223-25875302 in the Zebra Finch genome, mean *F*_ST_ = 0.75), chr 20 (1981369, mean *F*_ST_ = 0.90), and chr Z (66226657, *F*_ST_ = 0.82). Such genomic architecture of divergence revealed by GBS data is consistent with the pattern from the WGS data (Figure S3), where the genome is largely undifferentiated except a few peaks at chr 1A, 4, 5, 20, and Z (Figure S3G).

**Figure 2.**
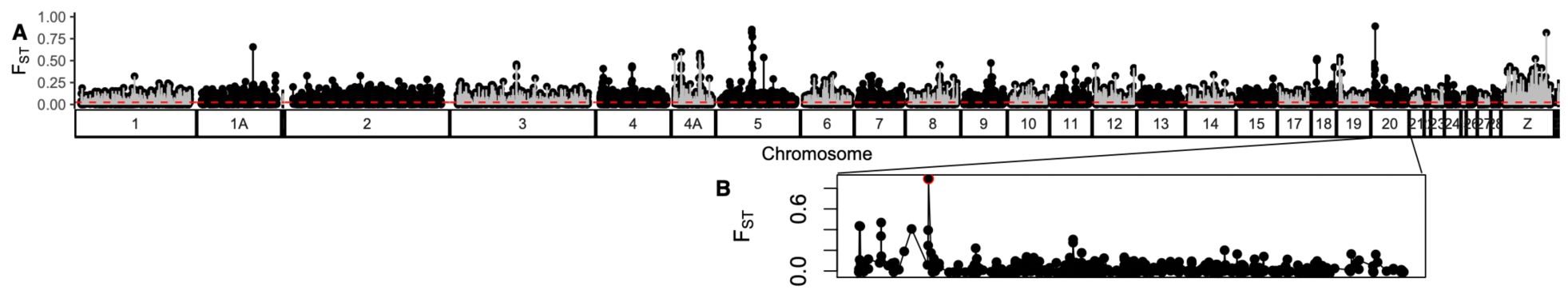
*F*_ST_ (between inland *townsendi* and *occidentalis*) scan is consistent with the isolation with gene flow model in which a few regions are under selection (divergent selection or selection against hybrids) while the rest of the genome is similar between species (Weir Cockerham Weighted average *F_ST_* is indicated by the horizontal red dotted line) due to gene flow. A, *F*_ST_ genomic scan of the genome revealed an island of differentiation around the highest peak in chromosome 20 (B, zoom in chromosome 20).

The *F_ST_* peak on chr 20 (Figure 2B) from the GBS data is within an intron of the gene RALY, which encodes heterogeneous nuclear ribonucleoprotein, the cofactor for cholesterol biosynthetic genes [69]. We hereafter refer to this SNP as the RALY SNP. This RALY SNP demonstrated the highest *F*_ST_ (0.90) inside an “island” of relatively high *F*_ST_ on chr 20 between *occidentalis* and *townsendi* parental populations across the genome (Figure 2B). The pure *occidentalis* population mostly contains GG homozygotes (referred to hereafter as OO), while the *townsendi* population contains CC homozygotes (referred to as TT). In the WGS data (Figure S4), which has much higher density of markers across the genome than the GBS dataset, this ‘island’ of differentiation contains pigmentation genes ASIP, EIF2S2, and RALY (Figure S4BC), thus we refer to it hereafter as the “ASIP-RALY” block. The RALY SNP revealed by the GBS data (Figure 2B) conveniently becomes the marker representing this ASIP-RALY divergent gene block surveyed across a large number of individuals (Figure S4BC).

Admixture mapping showed a strong pattern of few loci associated with divergent plumage traits in the hybrid zone. In particular, the colors of the crown, cheek, and the breast were each very strongly associated with the same RALY SNP (on chr 20) that was the outlier in the *F*_ST_ analysis (crown: *χ^2^* = 57.89, r^2^ = 0.57, *p* = 5.95 × 10^−15^, Figure 3A, Figure 4AD; cheek: *χ^2^* = 44.76, r^2^ = 0.55, *p* = 2.23 × 10^−11^, Figure 3C, Figure 4BE; breast: *χ^2^* = 28.47, r^2^= 0.57, *p* = 9.53 × 10^−8^, Figure 3D, Figure 4CF). The RALY SNP shows a partial dominance pattern of the G allele for the three plumage traits, with G/C heterozygotes tending to have similar phenotypes as G/G homozygotes, although there appears to be some additivity as well (Figure 4). The RALY gene (Figure 3B) is associated with yellow pigmentation in mice and quail [70, 71], and is adjacent to two other pigmentation genes -- ASIP (115941 bases away) and EIF2S2 (30223 bases away) (Figure 3B, S4C).

**Figure 3.**
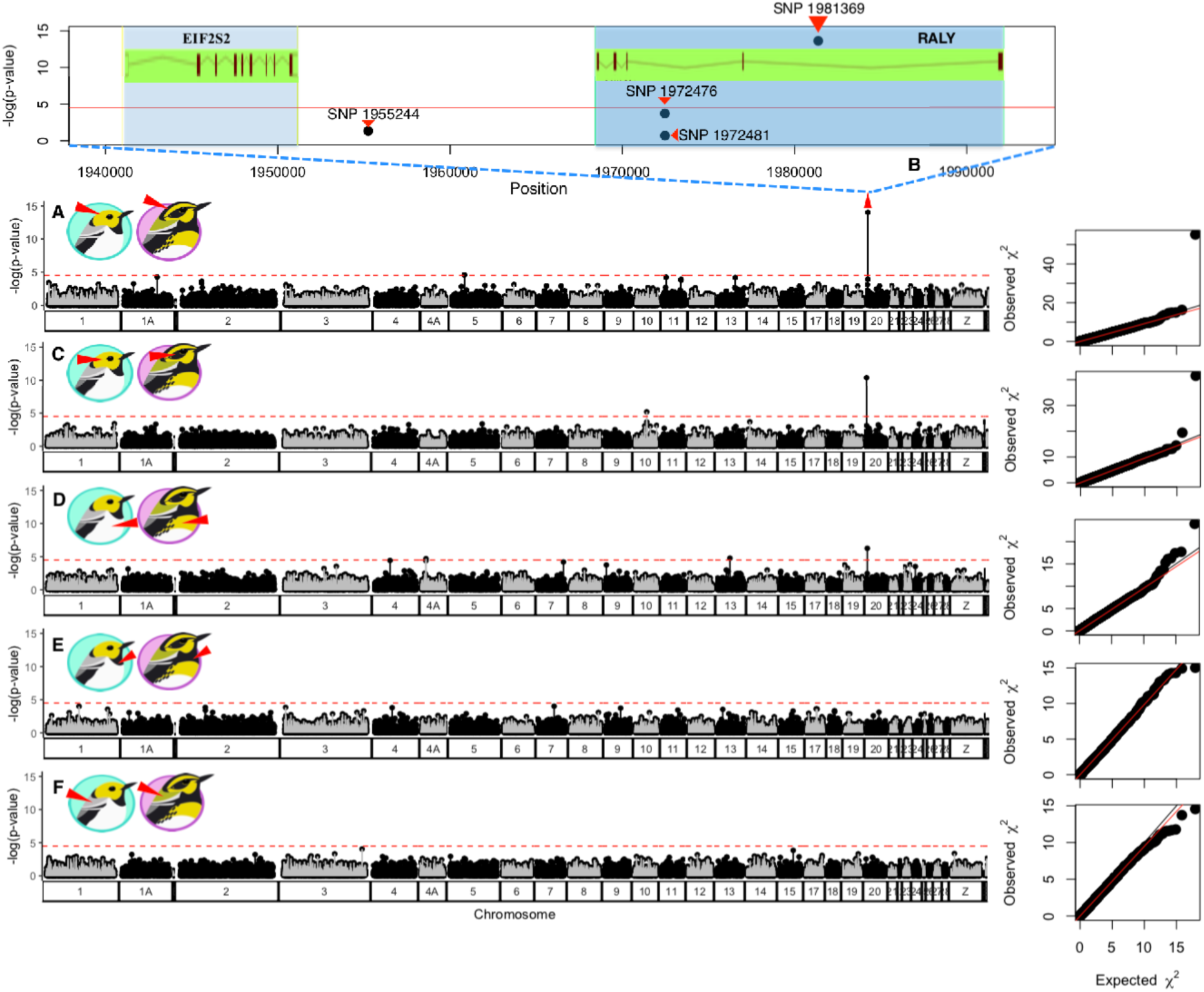
Admixture mapping revealed a SNP inside the RALY gene in chromosome 20 at position 1981369 that was significantly associated with crown (**A**, **B**) and cheek darkening (**C**), as well as the intensity of yellow on the breast (**D**), but not with bib size (**E**) nor with the color of the back (**F**). **A-F**, genomic scan of inflation factor-corrected –log (*p*-value) of genotype-phenotype association tests. The horizontal red lines represent the Bonferroni-corrected critical threshold. The plumage trait tested in each scan is indicated by the red triangle in the cartoons. **A-C**, there was a strong peak at chromosome 20 position 1981369 inside the RALY gene that indicates a pleiotropic effect on crown, cheek, and breast coloration. **B**, genetic map of around position 1981369 on chromosome 20 and its nearby SNPs in this study. The green boxes represent the stretches of the protein-coding genes, EIF2S2 and RALY, with the exons depicted as the red vertical bars connected by introns (red lines). **E-F**, no significant SNP was detected that is associated with bib size or greenish back.

**Figure 4.**
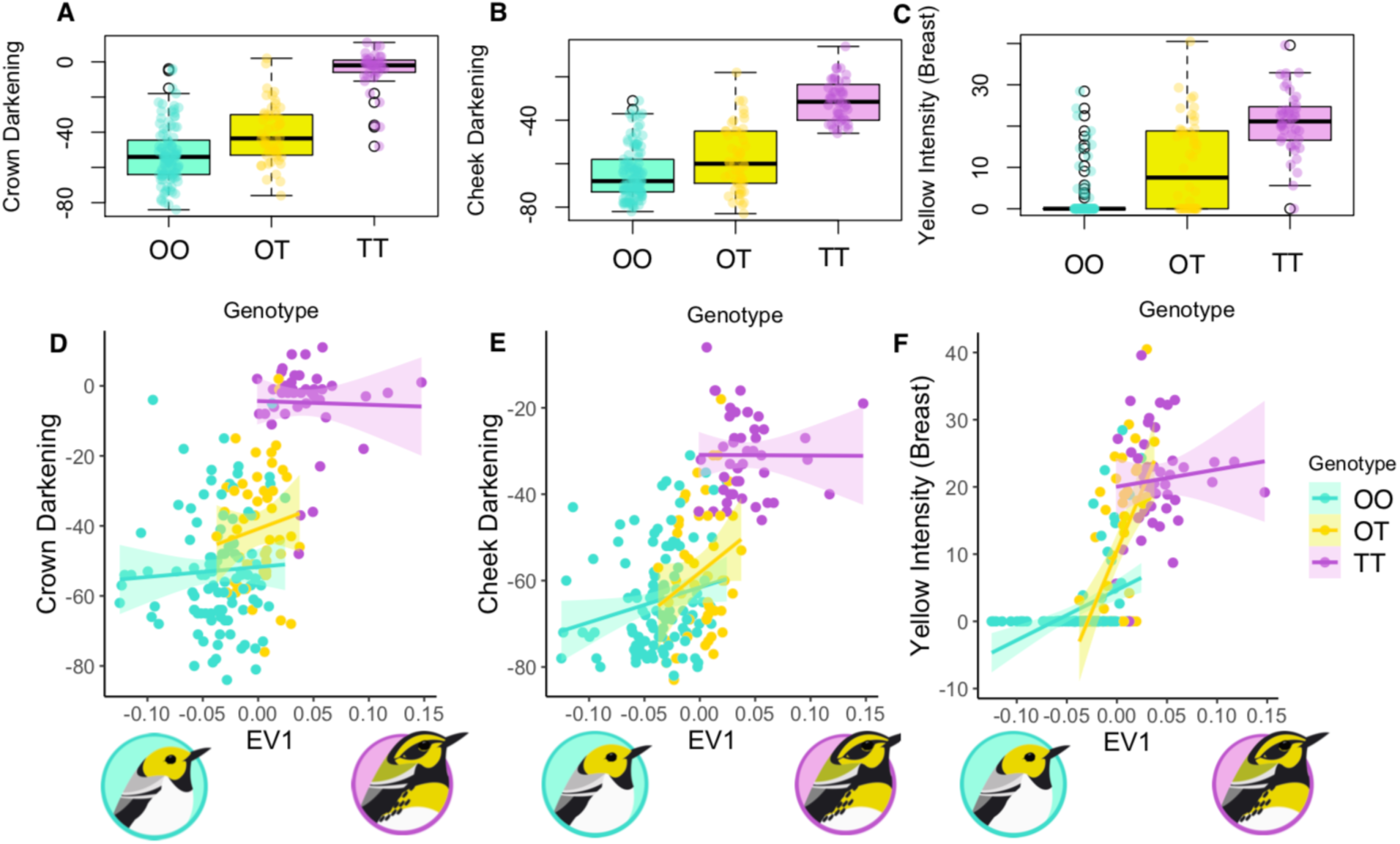
Association of the RALY SNP on crown (**A**, **D**), cheek (**B**, **E**), and breast (**C**, **F**) coloration, in which the pure *occidentalis* genotypes is denoted as “OO”, pure *townsendi* genotype is denoted as “TT”, and heterozygotes as “OT”. **D**-**F**, The associations of the RALY genotype and cheek darkening among individuals are significant (*p* < 10^-7^) after accounting for the underlying genomic ancestry. The data are consistent with a combination of dominance and additive effects on the three plumage traits.

A low-resolution sequencing approach such as GBS, which sequences less than 1% of the genome, would be unlikely to detect a SNP that is directly causal for phenotypic variation. It is more likely that the RALY SNP is closely physically linked to the causal DNA variant (SNP or some other type of variant) for the phenotypic differences. It is also possible that multiple linked SNPs in this region are responsible for the variation in different phenotypic traits, such that the apparent pleiotropic effect (i.e., strong association with three plumage traits) of the RALY SNP might be due to separate causal genes that are closely linked. The continuous divergence of the ASIP-RALY gene block between allopatric species in the WGS dataset suggests that the ASIP-RALY gene linkage block might be involved in plumage color divergence as a whole (Figure S4).

Three additional SNPs were found to be significantly associated with the intensity of yellow on the breast (Figure 3D). Two are on chr 4A at nucleotide location 5588139 and 5588235, both inside an ortholog of the mammal Immunoglobulin Binding Protein 1 (IGBP1) gene. One is at nucleotide location 6623798 on chr 13, inside gene Annexin A6, which codes for a phospholipid binding protein, Annexin VI [72]. We did not detect SNPs that significantly explained variation in either bib size or green coloration on the back (Figure 3EF). The RALY SNP explains 57% of variation in crown (Figure 4AD), 52% of the variation in cheek (Figure 4B, E), and 43% variation in breast coloration (Figure 4CF).

We found evidence that the ASIP-RALY region is under divergent selection, as it is the most extreme *F_ST_* outlier, while most of the genome is not very differentiated (average = 0.03) (Figure 5D). In addition, the RALY SNP (representing the ASIP-RALY gene block) demonstrated a spatial cline that was stable in location over two decades (Figure 5A). The cline center was at 1216.35 ± 4.0 (SE) km in 1987-94 (Figure 5A, blue curve), and did not significantly shift in 2015-16 sampling (Figure 5A, yellow curve), occurring at 1213.140 ± 2.2 (SE) km. Furthermore, the width of the RALY cline stayed narrower (95% CI of *w^2^_2015-16_* -*w^2^_1987-94_*: - 39004.70 to 1035.13 km^2^) than under a model of neutral diffusion (*w^2^_2015-16_* -*w^2^_1987-94_* = 7540 km^2^) (Figure 5B, see [46] for explanation of method).

**Figure 5.**
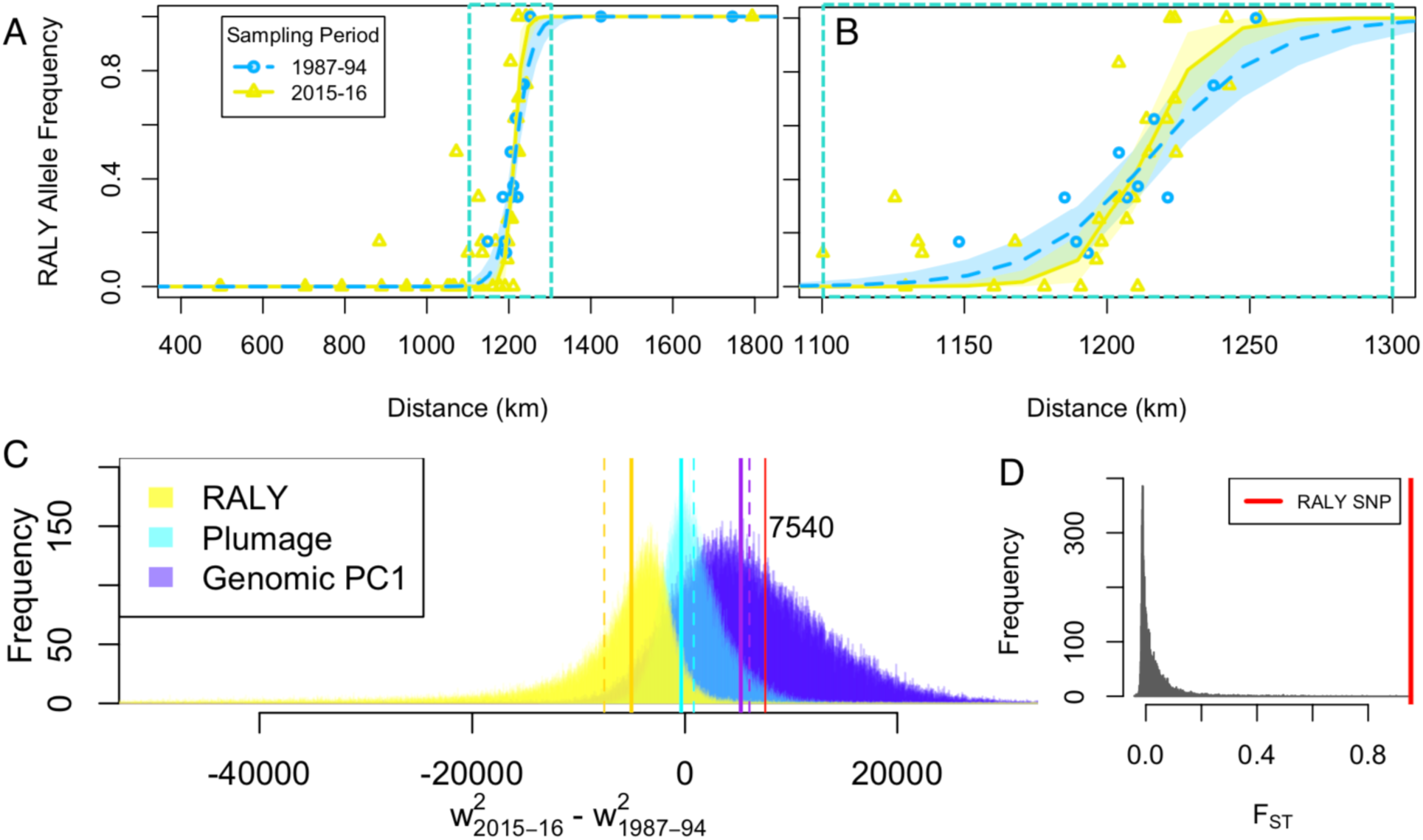
Evidence of selection on RALY locus or its linked region. The RALY SNP shows stable clines (sampled in 1987-94 and 2015-16) across the hybrid zone, which are extremely narrow (**A, B**, zoom in around 1100 to 1300 km), and has become significantly narrower in the recent sampling (**C**). The bootstrap distribution of *w^2^_2015-16_* -*w^2^_1987-94_* (the change of squared cline width between sampling periods) of the RALY cline is less than expected under neutral diffusion, suggesting selection has been maintaining narrow clines of genomic PC1, plumage, and RALY SNP (**C**). The selection at RALY (greater deviation from neutral expectation than plumage and genomic PC1) might indirectly maintain the narrow plumage and genomic cline (**C**). The vertical lines are *w^2^_2015-16_* -*w^2^_1987-94_* values (solid line: sample estimates; dotted line: bootstrap mean values) respectively for RALY cline (yellow), plumage cline (turquoise), and genomic PC1 cline (purple). (**D)**, RALY is the extreme outlier in the distribution of *F*_ST_ between inland *townsendi* and *occidentalis* across the genome (RALY SNP *F*_ST_ value depicted by the red line).

## Discussion

We observed clear but narrow genomic ‘islands of differentiation’ between Townsend’s and Hermit Warblers on chr 1A, 5, 20, and Z (Figure 2). This agrees with the predicted pattern of speciation in the face of gene flow: as most of the genome remains homogenized due to gene flow and lack strong selection, restricted targets of divergent selection have become strongly differentiated [3,7,11,73].

The RALY SNP (chr 20) within the ASIP-RALY gene block represents one of the ‘islands of differentiation’ and is strongly associated with key divergent traits: cheek, crown, and breast coloration. The consistently narrow geographical RALY cline in the hybrid zone over several decades, despite ongoing gene flow [46], further implicates selection (divergent selection and/or selection against hybrids) around the RALY SNP as contributor to reproductive isolation between these young sister species. The *w^2^_2015-16_* -*w^2^_1987-94_* estimate at the RALY cline was less than the plumage cline and the genomic PC1 cline (Figure 5B), suggesting that a narrow region centered at the RALY locus might be the direct target of selection maintaining the stable plumage and genomic clines [46]. Such selection may further extend genomic differentiation by elevating the differentiation at linked sites.

Although there is signature of genetic hitchhiking that underpins extended genetic divergence around the ASIP-RALY targets of selection (Figure S5), it seems that the plumage pigmentation effect is more narrowly underpinned by ASIP-RALY region. In the hybrid zone, the recombination over multiple generations can break down the hitchhiking gene blocks (that tend to co-segregate with the trait-determining loci in allopatric populations), revealing specific genetic underpinnings of the differentiated phenotypes. The genetic underpinning of the plumage trait should be very close to the narrow region around the RALY SNP.

The ‘divergence with gene flow’ pattern, in which closely related taxa hybridize extensively and have little differentiation across much of the genome, along with evidence of a narrow and highly differentiated genetic region that has large effect on divergent traits suggests a simple genomic architecture of divergence. Such a strong target of selection could effectively counteract gene flow and maintain a species boundary at secondary contact, allowing speciation to progress [4,74,75]. There may also be weak multilocus divergent selection on a large set of loci, yet such weak polygenic selection is inherently challenging to detect.

### Pleiotropy

This ASIP-RALY block underlying multiple plumage patches could provide a strong pleiotropic mechanism (one region that affects multiple phenotypes [76]), linking divergent selection and possibly reproductive isolation. The RALY SNP is associated with a suite of melanin- and carotenoid-related plumage patches (crown, cheek, and breast coloration) that differ diagnostically between species (Figure 3) and are likely under some form of selection. Previous admixture mapping in *Setophaga coronata auduboni*/*S. c. coronata* revealed SCARF2 on chr 15 that is associated with multiple carotenoid and melanin traits [77]. Intriguingly, in this sister pair within the same genus, drastically different color regions are involved in carotenoid versus melanin plumage divergence. Such distinct pleiotropic genetic underpinnings of plumage divergence may facilitate rapid speciation in *Setophaga* [78].

Melanin and carotenoid plumage traits are often involved in mate-choice and male-male competition [79–82]. If carotenoid-related plumage traits are targets of female mate-choice or male-male competition for mating opportunities in *Setophaga* warblers, strong selection on pleiotropic genes can play a powerful role in limiting mating between species and facilitating speciation. Pleiotropy is a more effective mechanism linking selection to reproductive isolation than linkage or polygenic inheritance, although genetic evidence for pleiotropic genes that influence both selected traits and reproductive isolation has been scarce (see review [83]). Among the limited existing examples, the classic cases involve coloration: the wingless gene affecting reproductive isolation and wing coloration in *Heliconius* butterflies [19], and the YUP locus affects pollinator isolation and flower coloration in monkey flowers [10]. RALY could be an additional example of a speciation gene. The stable and narrow cline of RALY SNP and extreme differentiation at the RALY locus imply divergent selection and/or selection against hybrids at this locus, which suggests that RALY could be associated with reproductive isolation in this system. Future study is needed to validate the possibility by examining the role of RALY-ASIP-regulated plumage signals in mate preference or male-male competition.

### Dominance

Our results support the prediction that Rohwer and Wood [43] made two decades ago that the cheek coloration in hybrids is underpinned by a single dominant locus. Indeed, heterozygotes of the RALY SNP exhibit similar yellow cheeks as the homozygous GG individuals (i.e., yellow cheek), which were found predominantly in the *occidentalis* population (Figure 4D-F). Dominance in the signal trait might reduce gene flow if it is in the opposite direction of dominance in the receiver trait (i.e., heterozygous receivers are more sensitive to *townsendi* signals), due to ‘opposing genetic dominance’ in receiver and signaler traits [83, 84]. This is because the F1 hybrids (heterozygous for both signaling and receiving) might for example, demonstrate *occidentalis* signal and preference for *townsendi*, thus cannot mate with other F1 hybrids. Therefore the *occidentalis* RALY dominance effect on the cheek signal could suppress gene flow if it is opposed by a *townsendi* dominant effect on receiving trait.

### RALY SNP

The RALY SNP is at the peak of divergence at allopatry (Figure 2B) and is tightly associated with plumage variation in the hybrid zone, although its neighboring SNPs are not significantly associated with plumage traits (after controlling for population substructure) (Figure 3, 6C). The roughly 180kb extended divergence around RALY in WGS data (Figure S2, S4) suggests the entire color gene block (ASIP-RALY) is involved in divergence.

The RALY gene encodes for heterogeneous nuclear ribonucleoprotein, the RNA binding protein that is known to regulate the expression of its downstream gene that codes for agouti signaling protein (ASIP) [70]. Deletion of RALY in Japanese quails and mice leads to novel transcripts of ASIP, which cause a yellow skin phenotype called “lethal yellow” [70, 71]. In mice, a lethal yellow mutation (Ay) in ASIP exhibits dominant pleiotropic effects on other traits in addition to skin coloration, which includes obesity, diabetic condition, etc., that are unrelated to the ASIP gene [71]. ASIP itself is well known for its influence on skin pigment by binding to melanocortin receptors and competitively excluding its agonists, preventing black/brown pigmentation [85, 86]. In birds, ASIP has been repetitively involved in genomic divergence between closely-related species in *Vermivora* [26], *Sporophila* [87], and *Munia* [88], in a mendelian inheritance pattern with the melanic form being the recessive phenotype. Genetic variants of RALY in warblers could result in differential expression ASIP leading to variations in carotenoid and melanin patterning. The pleiotropic dominance effect of the RALY SNP we identified in this natural hybrid zone is consistent with these known effects of RALY mutants in the lab. Future study should investigate the functional interaction of RALY and ASIP in plumage divergence, and whether there is any convergent mutation in ASIP between *Vermivora* and *Setophaga*.

On top of the strong pleiotropic ASIP-RALY association with plumage, there could be other regions of the genomes contributing to further refinement of species-specific coloration in this system. Specifically, other significant SNPs on chr 4A and 13 are associated with breast coloration. In addition, there might be narrow but important genetic regions that this GBS data did not cover. However, this is unlikely because the WGS data did not reveal additional differentiation peaks.

## Conclusion

We identified a few narrow and dispersed genomic regions of differentiation between two hybridizing *Setophaga* warblers, consistent with divergent selection in the face of gene flow. A highly divergent peak spans among three closely linked pigmentation genes: ASIP, EIF2S2, and RALY. Admixture mapping within a hybrid zone between these species showed that this SNP was strongly associated with three distinct plumage traits that distinguish the species, revealing a major-effect region related to speciation. In contrast, we did not find genetic regions significantly associated with the other two species-diagnostic traits, implying many genes of small effects.

The extensive similarity in the rest of the genome raises a question regarding the distinctiveness of *townsendi* and *occidentalis* and their future. Key to this question is whether these regions of differentiation represent sufficient reproductive isolation that will maintain separation for further differentiation to build up. The genetic divergence of the ASIP-RALY gene region between allopatric populations and the stable narrow geographical cline of this region suggest that selection maintains such difference between species. Future studies should track the genomic differentiation in linkage disequilibrium with the ASIP-RALY region and investigate the mechanism of apparent selection. Overall, these results show that avian sister species can be very similar across the great majority of their genomes, sometimes differing primarily at narrow genomic regions under selection maintaining phenotypic differences between species.

## Acknowledgement

Ruth Midgley and Else Mikkelsen provided excellent assistance in the field and lab for this study. We thank Chris Wood (Burke Museum) for help relocating the historical sites and accessing the historical plumage samples, and Sharon Birks (Burke Museum) for accessing tissue samples for sequencing. Sally Otto, Ailene MacPherson, Dolph Schluter, Loren Rieseberg, Armando Geraldes, Tom Booker, Toby Bradshaw, Molly Schumer, and members of the Irwin Lab provided helpful discussion. We thank research funding provided by the Natural Sciences and Engineering Research Council of Canada (grants 311931-2012, RGPIN-2017-03919 and RGPAS-2017-507830 to DEI; and PGS D 331015731 to SW), a Werner and Hildegard Hesse Research Award in Ornithology and a UBC Four Year Doctoral Fellowship to SW. Finally, we thank Environment Canada, U. S. Geological Survey, Departments of Fish & Wildlife of Washington, Oregon, California, Idaho, and Montana, and the UBC Animal Care Committee for providing research permits.

## Supplementary Information

We then investigated whether selection at the RALY marker representing ASIP-RALY gene block has led genetic hitchhiking of its flanking regions facilitating further differentiation around this target of selection. We tested whether the bootstrap (for 10,000 iterations) difference of *F_ST_* between the candidate genetic block (300kb flanking region of the ASIP-RALY genetic block, excluding the RALY locus itself, colored in yellow Figure S5A) versus the rest of chr20 (colored in light blue Figure S5A) is significantly greater than 0. Indeed, we found the signature of hitchhiking, as ASIP-RALY candidate genetic block demonstrated significantly greater *F_ST_* than the rest of the regions in chr20 (Figure S5AB, grey distribution of difference is greater than 0, 95% CI: 0.107-0.109, t_9999_ = 254.31, *p* < 10^-15^).

**Figure S1.**
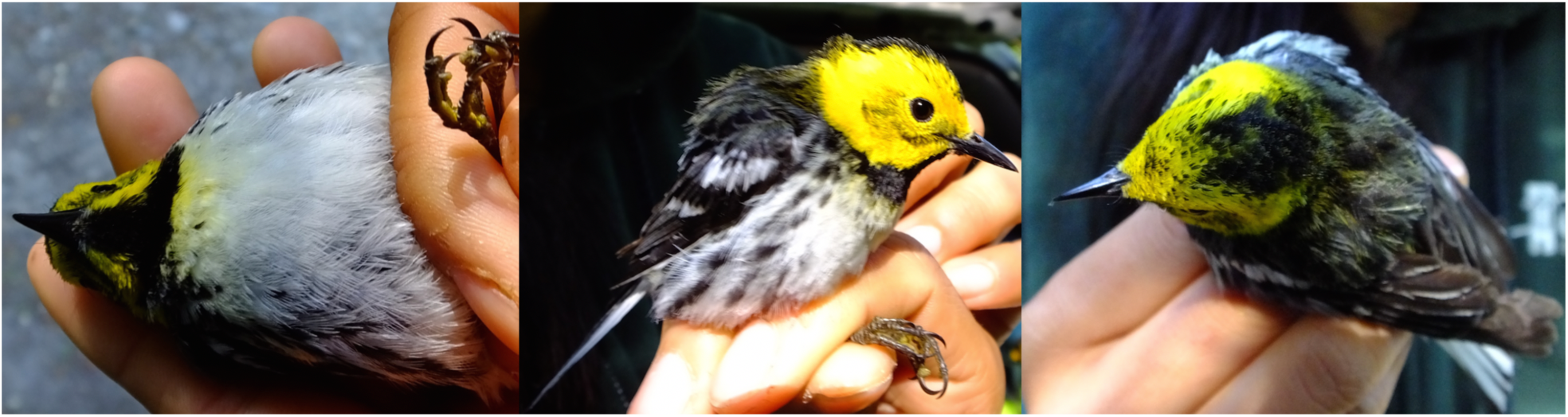
Field photos of a hybrid male showing three different angles: 1) frontal with head tilted up showing throat badge and breast measurements), 2) profile showing the cheek, and 3) from above showing the crown.

**Figure S2.**
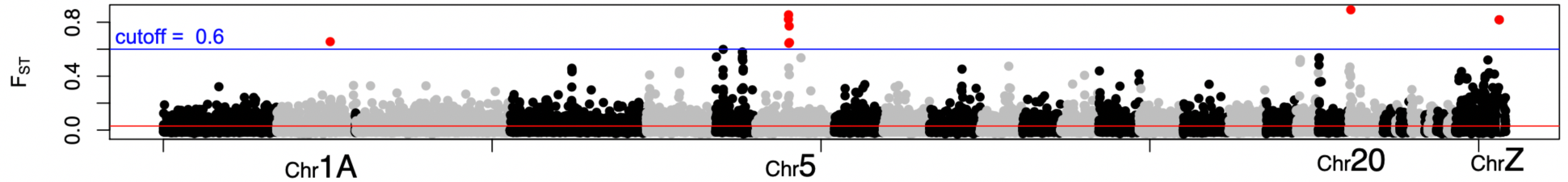
SNPs with *F_ST_* > 0.6 are colored in red. High *F_ST_* SNPS are distributed on chr1A, 5, 20, and Z.

**Figure S3.**
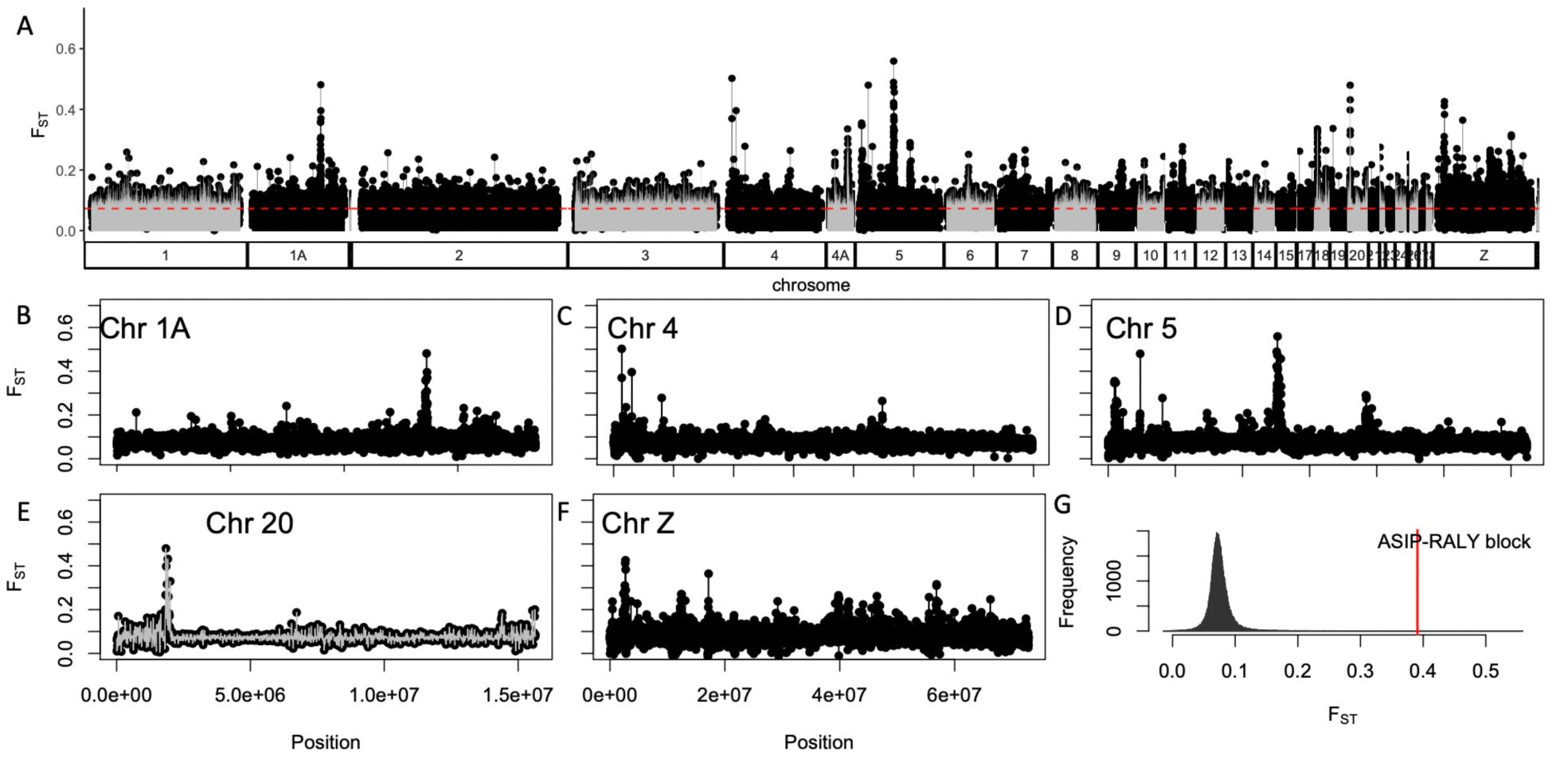
Weir Cockerham *F_ST_* scan of with the WGS data, in which each dot represents a 10kb non-overlapping window (A), where peaks were found on chromosome 1A (B), 4 (C), 5 (D), 20 (E), and Z (F). G, the ASIP-RALY gene block demonstrates extremely high *F_ST_* relative to the rest of the genome.

**Figure S4.**
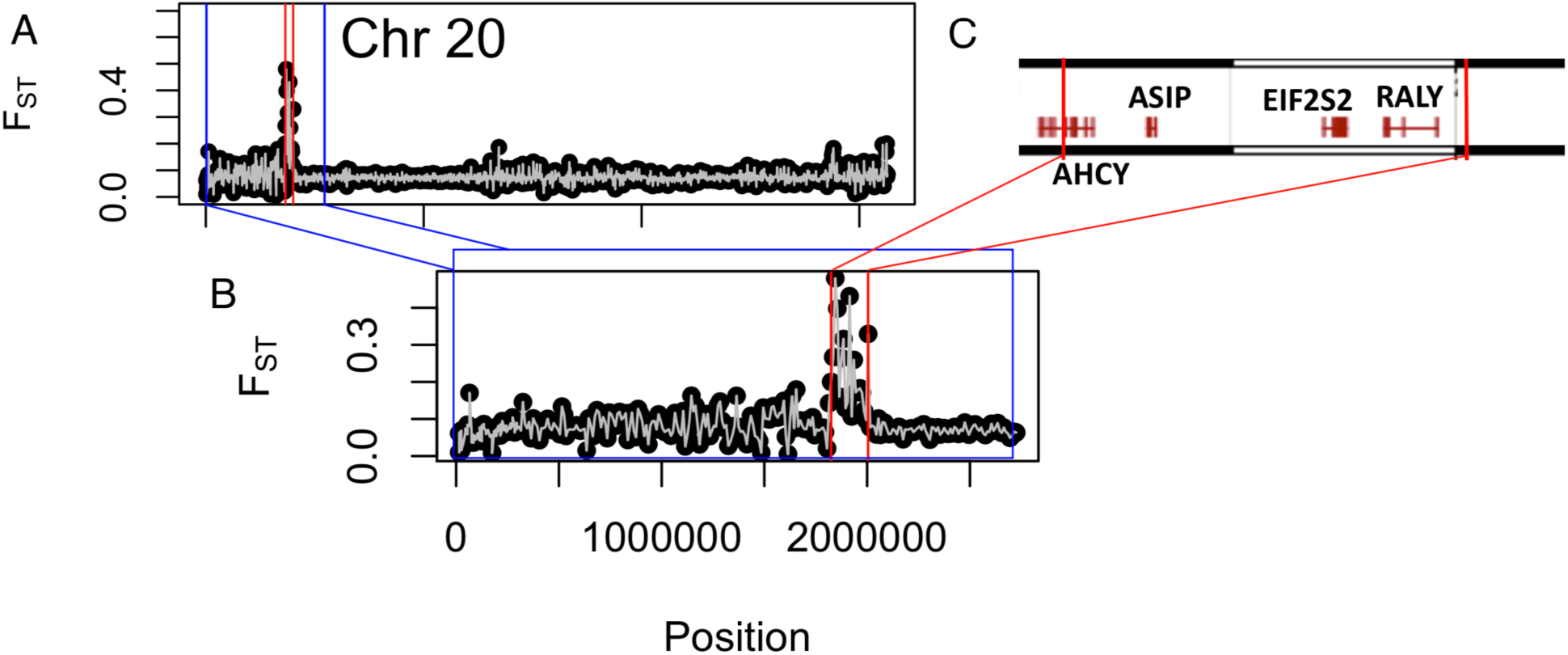
The ***F_ST_*** peak on chromosome 20 (A) resides in the ASIP-RALY gene block (B, C). C, the position of the protein coding genes relative to the peak (bounded by the red vertical lines). C, within the genes, the vertical strikes are the coding regions flanked by the non-coding regions (horizontal lines).

**Figure S5.**
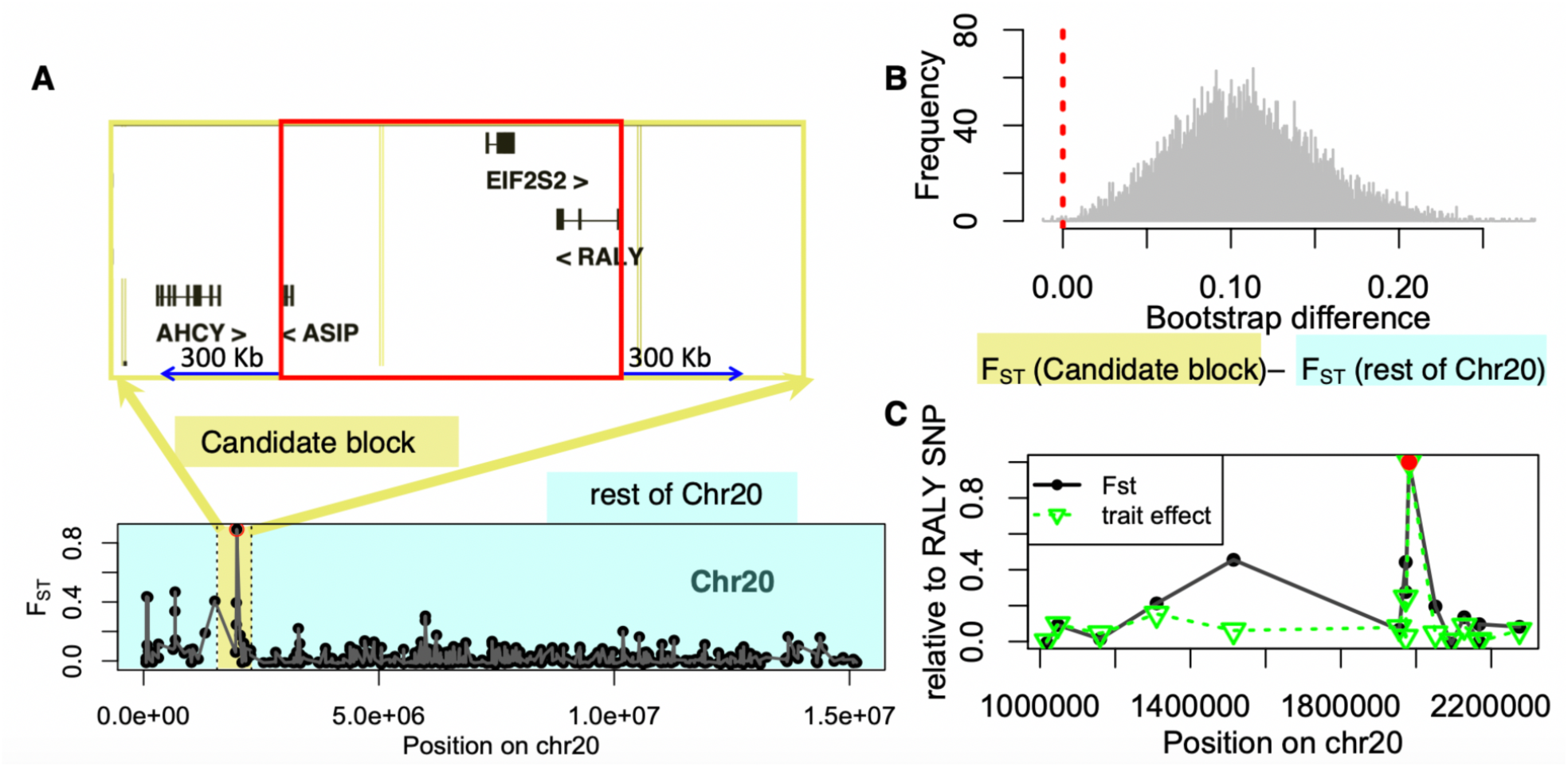
There is signature of hitchhiking around the RALY peak (elevated *F_ST_* around RALY), while the effect on plumage coloration is more tightly associated with RALY. Thus the potential causal locus of this trait should be very close to RALY. **A, B** signature of hitchhiking: the candidate gene block (the pigmentation gene block and its 300kb flanking region excluding the RALY locus) has higher *F_ST_* than regions in the rest of the chr20 (B, the bootstrap difference of *F_ST_* is significantly greater than 0, indicated by the vertical dotted line, 95% CI: 0.107-0.109). **C**, in comparison to *F_ST_*, the trait effect is more specific to the RALY peak: there is a steeper decay of scaled trait effect size (relative to RALY peak) than the scaled differentiation (relative *F_ST_* to RALY peak).

**Table S1.** Allele frequencies of SNPs between RALY and ASIP in hybrid zone and parental zones. The causal genetic mechanism for plumage coloration might be narrower than the ASIP-RALY linkage block around the RALY SNP. The RALY SNP is in physical proximity to the two other pigmentation genes mentioned above, although 3 other SNPs (1955244, 1972476, and 1972481) that are 8888-26125 bp away from RALY SNP (Table S2; Figure 3B), physically closer to ASIP and EIF2S2, did not show association with phenotype (Figure 3B). Two of these SNPs showed low minor allele frequencies (0.05-0.09) in the hybrid zone, and thus are not expected to be highly associated with trait variation. However, the fact that SNP 1973476 (closer to the other genes than the candidate RALY SNP) demonstrated similar minor allele frequency as the significant RALY SNP and was not significantly associated with plumage coloration highlights the importance of the region around position 1981369 inside the RALY gene. Either way, the RALY SNP represents the ASIP-RALY gene block and appears to have an effect on multiple species-diagnostic coloration traits.

| POS | allele1:freq | allele2:freq |
| --- | --- | --- |
| Hybrid zone |  |  |
| 1955244 | T:0.946 | A:0.054 |
| 1972476 | A:0.72 | C:0.28 |
| 1972481 | T:0.917 | C:0.083 |
| 1981369 | C:0.375 | G:0.625 |
| *townsendi* zone |  |  |
| 1955244 | T:1 | A:0 |
| 1972476 | A:0.5 | C:0.5 |
| 1972481 | T:0.656 | C:0.344 |
| 1981369 | C:1 | G:0 |
| *occidentalis* zone |  |  |
| 1955244 | T:0.95 | A:0.05 |
| 1972476 | A:0.944 | C:0.056 |
| 1972481 | T:0.972 | C:0.028 |
| 1981369 | C:0.048 | G:0.952 |

**Table S2.**
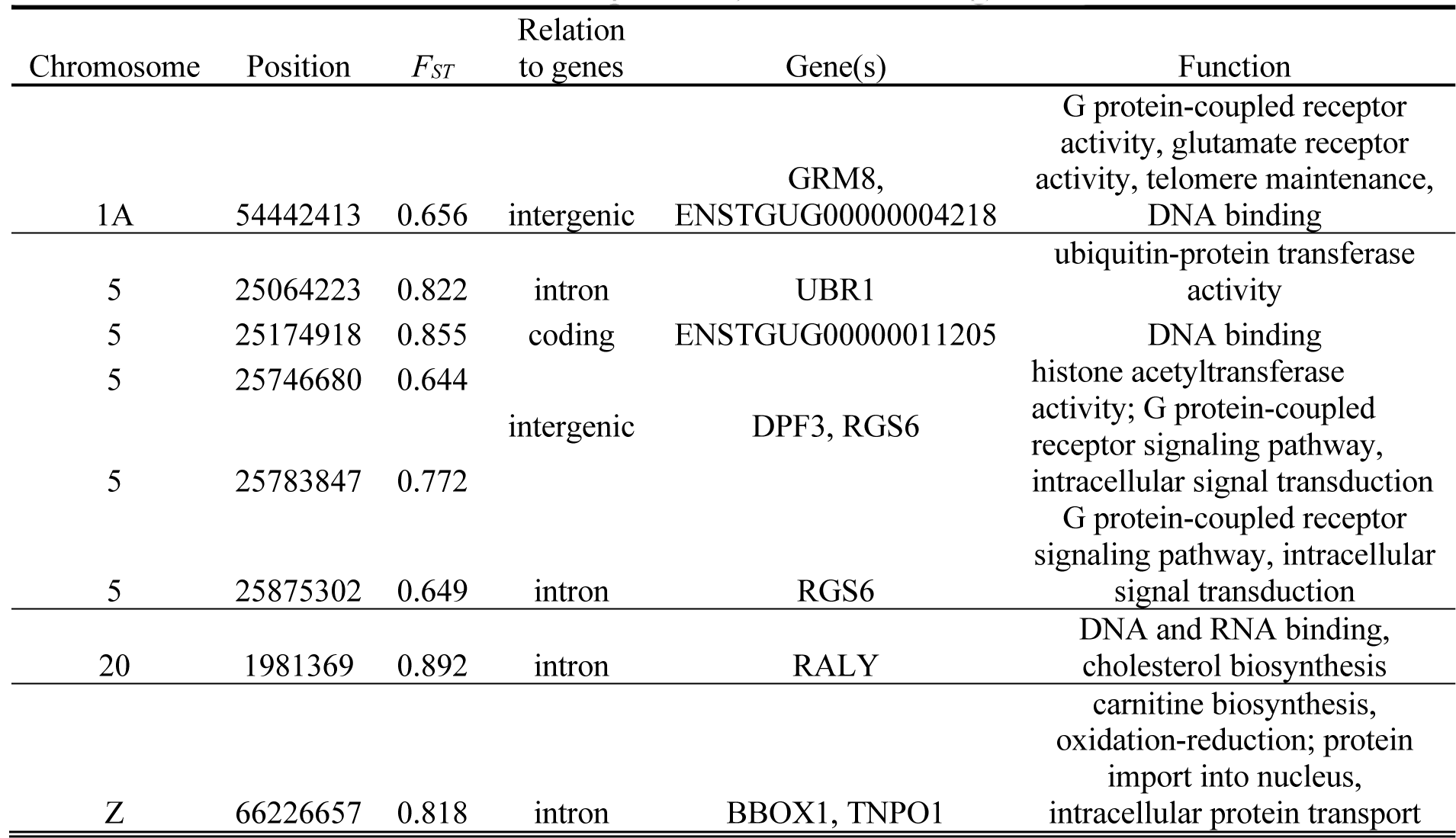
SNPs with *F_ST_* > 0.6 and their position, association to genes and molecular functions

